# ABI3 deletion in TgCRND8 mice is associated with reduced amyloid plaque pathology and altered glial response

**DOI:** 10.1101/2023.10.05.560956

**Authors:** Deniz Ghaffari, Jennifer Griffin, Peter St George-Hyslop

## Abstract

Recent genome-wide association studies (GWAS) have identified several rare coding variants in a cluster of immune-related genes that are significantly associated with Alzheimer’s disease. This includes the risk variant (S209F) in *ABI3*, a gene encoding the Abl Interactor family member 3 protein that is highly expressed in microglia, but little is known about its function in these cells and its association with AD. We have investigated the effects of ABI3 deletion on AD pathological features in the transgenic (Tg) CRND8 APP mouse model by immunofluorescence (IF) staining and confocal microscopy. We have observed that ABI3 expression is localized to microglia in the mouse brain and that the ABI3 immunoreactivity levels are increased in the TgCRND8 mice. The loss of ABI3 leads to a significant reduction of amyloid plaque numbers and size in the 9-month-old TgCRND8 mice. This is also accompanied by a reduced number of microglia clustering around the plaques. On the basis of these observations, we hypothesize that the loss of ABI3 may lead to changes in microglial behavior which directly or indirectly leads to reduced AD pathology in mice.

## Main

Alzheimer’s disease (AD) is the most common neurodegenerative disorder with distinct pathological hallmarks including the combined presence of aggregated extracellular Amyloid-beta plaques, intracellular neurofibrillary tangles, and glial activation, leading to chronic neuronal damage and progressive cognitive decline^1^. Currently, the innate immune system activation, in particular microglia, is considered a contributing factor to AD pathogenesis rather than a consequence of pathology^2^. This is largely due to the identification of several AD-associated risk genes with enriched expression levels in microglia and myeloid cells, through whole-exome sequencing and genome-wide associations studies (GWAS).

Amongst these genes is the recently identified gene *ABI3* which encodes the Abl Interactor family member 3 protein. The S209F variant of ABI3 (rs616338:p.Ser209Phe) is associated with increased risk of AD (OR = 1.43, p = 4.5 × 10 – 10, MAF = 0.008)^3^. ABI3 is the most recently discovered member of Abl Interactor (ABI) proteins, also including ABI1 and ABI2, that all possess a similar structure, containing a C-terminus Src homology 3 (SH3) domain and several serine-rich and proline-rich regions^4^. Similar to other ABI proteins, ABI3 is also believed to be involved in the actin cytoskeleton regulation by participating in the WASP family Verprolin homolog protein (WAVE) regulatory complexes (WRC), but its exact function in the cell remains unknown^5^. Outside of the central nervous system (CNS) ABI3 is highly expressed in the spleen, lymph nodes, and appendix. In the brain, the ABI3 transcript is present in microglia and at lower levels in non-myeloid cells including neurons^6^.

Following a thorough validation of the commercially available anti-ABI3 antibodies using ABI3 KO primary microglia as control cells, we detected ABI3 protein expression in Iba-1 positive microglia/macrophages in the mouse brain by immunofluorescence (IF) and the loss of ABI3 expression in the ABI3 knockout (^−/−^) mouse line (B6N(Cg)-Abi3tm1.1(KOMP)Vlcg/J) by western blotting and IF. It’s important to note that as reported by other groups, in the B6N(Cg)-Abi3tm1.1(KOMP)Vlcg/J mouse line the loss of ABI3 is accompanied by the ablation of the neighboring gene, *GNGT2*, therefore we will be referring to these mice as ABI3-GNGT2^−/−7^. Notably, the knockout of ABI3-GNGT2 led to a reduction of amyloid beta pathology and the number of plaque-associated microglia in the TgCRND8 APP mouse model, suggesting that ABI3 could modulate the risk of AD through regulating microglial response to amyloid pathology.

We tested the commercially acquired ABI3 antibody (rabbit anti-ABI3, abcam ab81152) for its sensitivity in selectively detecting ABI3 protein by western blotting and immunostaining in ABI3^−/−^ derived microglia lysates and brain tissue sections respectively. Previous reports have shown ABI3 expression in microglia and increased expression levels in AD brains, especially in the microglia surrounding the amyloid plaques^8^. In support of these observations, here we show that ABI3 immunoreactivity (red) is mainly localized to Iba-1 positive microglia/macrophages (green) in two hippocampal regions (Dentate Gyrus [**Figure 1.A**] and CA1 [**Figure 1.B**]) of 9-month-old C57/BI6 non-transgenic (NonTg) mouse brain, using IF.

**Figure 1:**
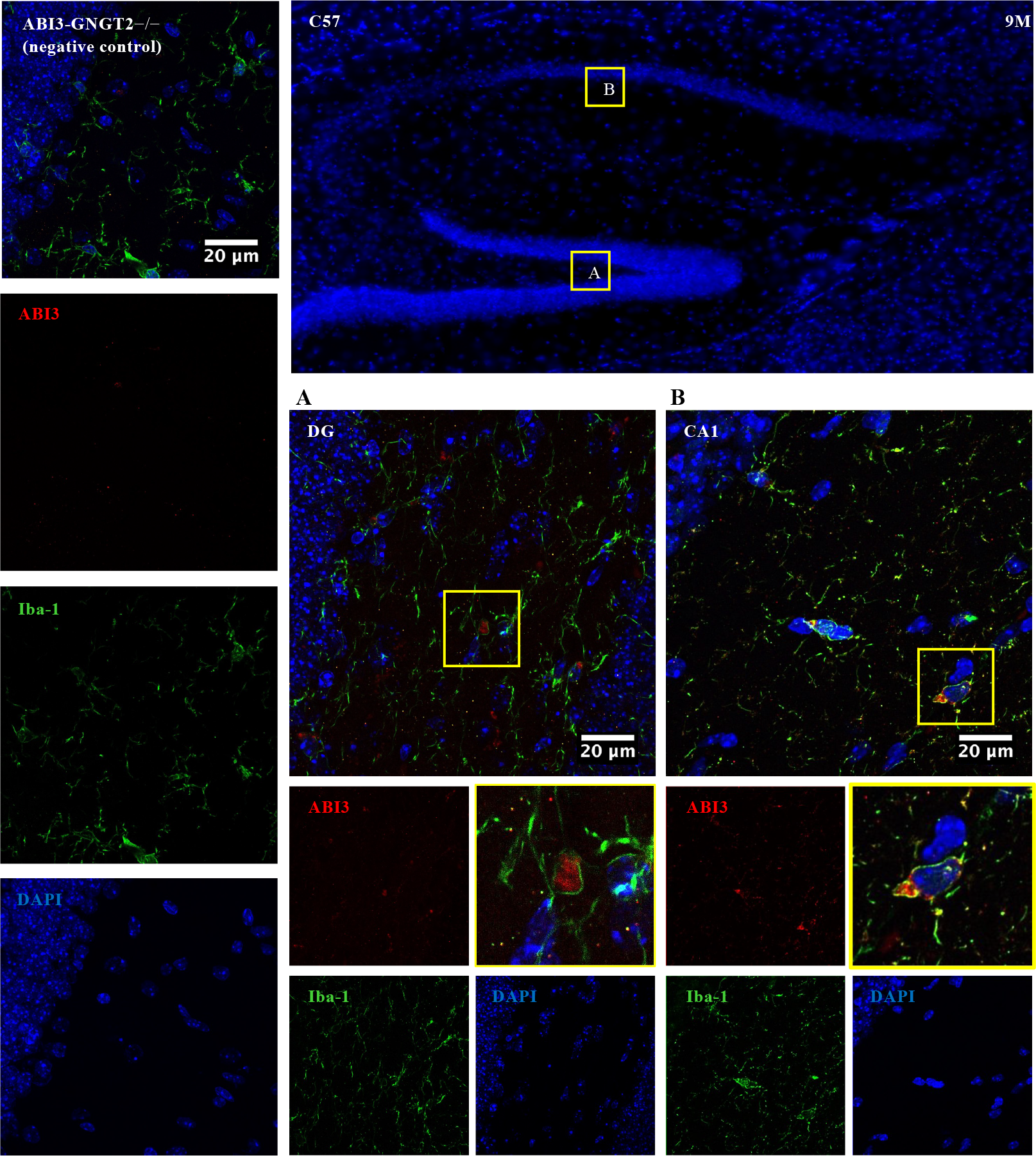
ABI3 immunoreactivity is observed in Iba-1 positive cells in C57 mouse brain tissue. Representative images of ABI3 (red) and Iba-1 (green)staining in the hippocampus of a 9-month-old C57 mouse brain. ABI3 expression was observed in Iba-1 positive cells in **A**. the Dentate gyrus (DG) and **B**. CA1 region of the hippocampus. Immunohistochemistry analysis of age-matched ABI3-GNGT2−/ −mouse brain tissue was performed in parallel as negative control where no ABI3 immunoreactivity was observed in Iba-1 positive microglia. N=10 mice (two sections from each mouse).

Prior research has also shown increased ABI3 gene expression in human primary monocytes following long-term (24 hours) immune stimulation by IFN-γ^9^. The ABI3 gene is located within a cluster of loci containing microglial genes that have been identified by GWAS, suggesting that they are components of a network of genes that modulate the risk of developing LOAD. Therefore, we were interested in investigating the changes in ABI3 expression during AD pathology and in TgCRND8 mice. For this purpose, we compared ABI3 immunoreactivity levels in TgCRND8 mice to age-matched non-transgenic C57 control mice (9-month-old) by IF. ABI3 immunoreactivity per cell in microglia is significantly increased in TgCRND8 mice relative to non-transgenic age-matched mice [**Figure 2**]. The increased ABI3 staining (red) was especially prominent in the microglia (green) surrounding the amyloid plaques (magenta) in the cortex and hippocampus of the mouse brain tissues studied.

**Figure 2:**
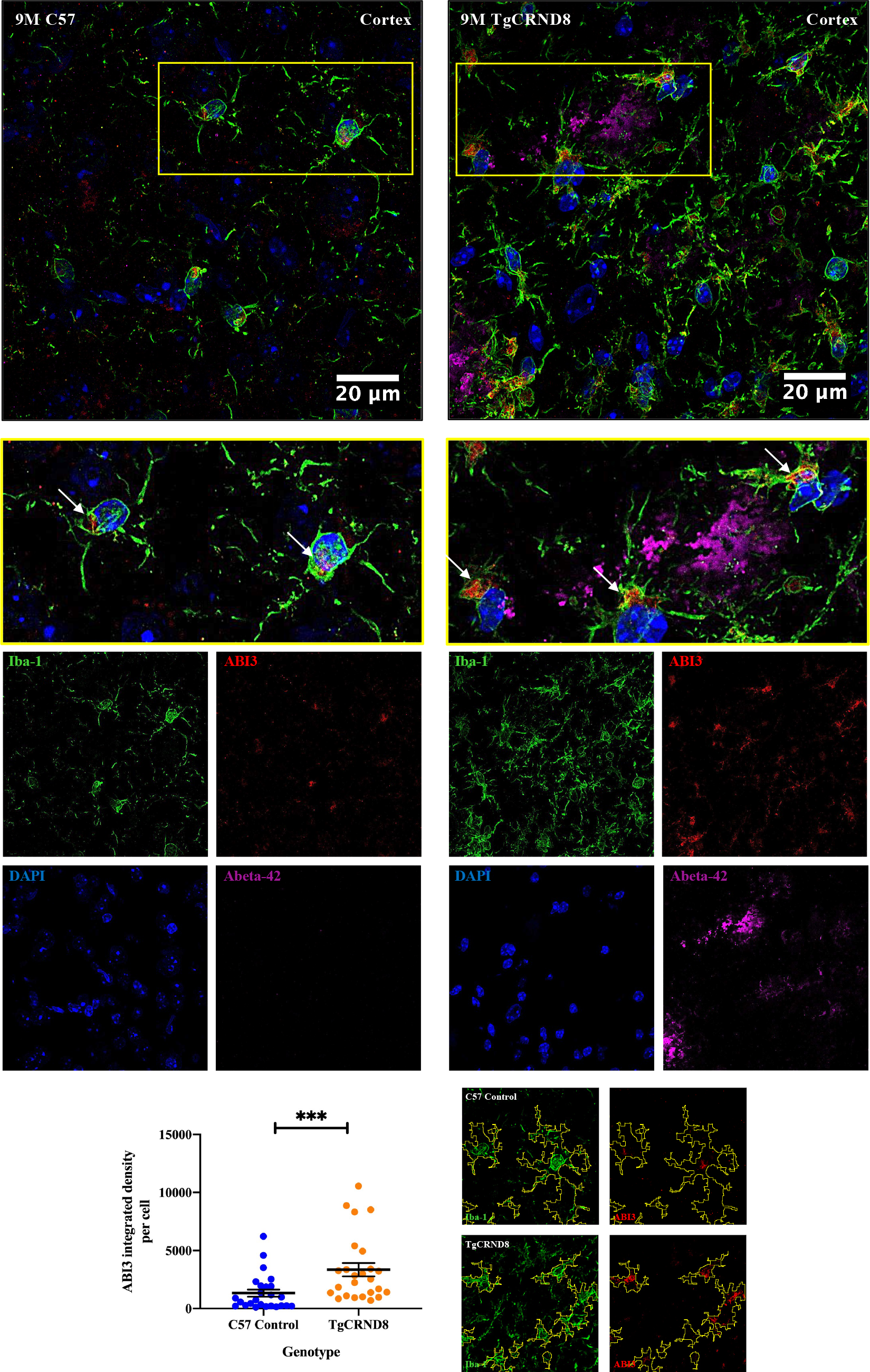
ABI3 immunoreactivity in microglia is increased in TgCRND8 mice. Representative images of ABI3 (red) and Iba-1 (green)staining in the cortex of 9-month-old C57 control (first row, left) and the microglia associated with amyloid-beta plaques (magenta) in TgCRND8 mouse brain tissues (first row, right). Arrows indicate Iba-1 (microglia) with ABI3 signal. N=10 (C57 9-month-old mice) and n=8 (TgCRND8 9-month-old mice) (two sections from each mouse). The statistical significance was determined using the unpaired t-test with Welch’s correction. (P value∼ 0.0004).

We examined if the increased ABI3 immunoreactivity in microglia was in relation to the microglia distance from amyloid plaques. Using IF and confocal microscopy we took several measurements of ABI3 integrated density in Iba-1 positive microglia in 9-month-old TgCRND8 mouse brain tissues in relation to the distance of the cell from the center of amyloid plaques, and plotted the results onto a scatter graph. We observed a moderate negative correlation between ABI3 integrated density levels and the distance of Iba-1 positive microglia from the plaque center, suggesting that ABI3 might be performing specific functions in the microglia populations that cluster around amyloid plaques [**Figure 3**].

**Figure 3:**
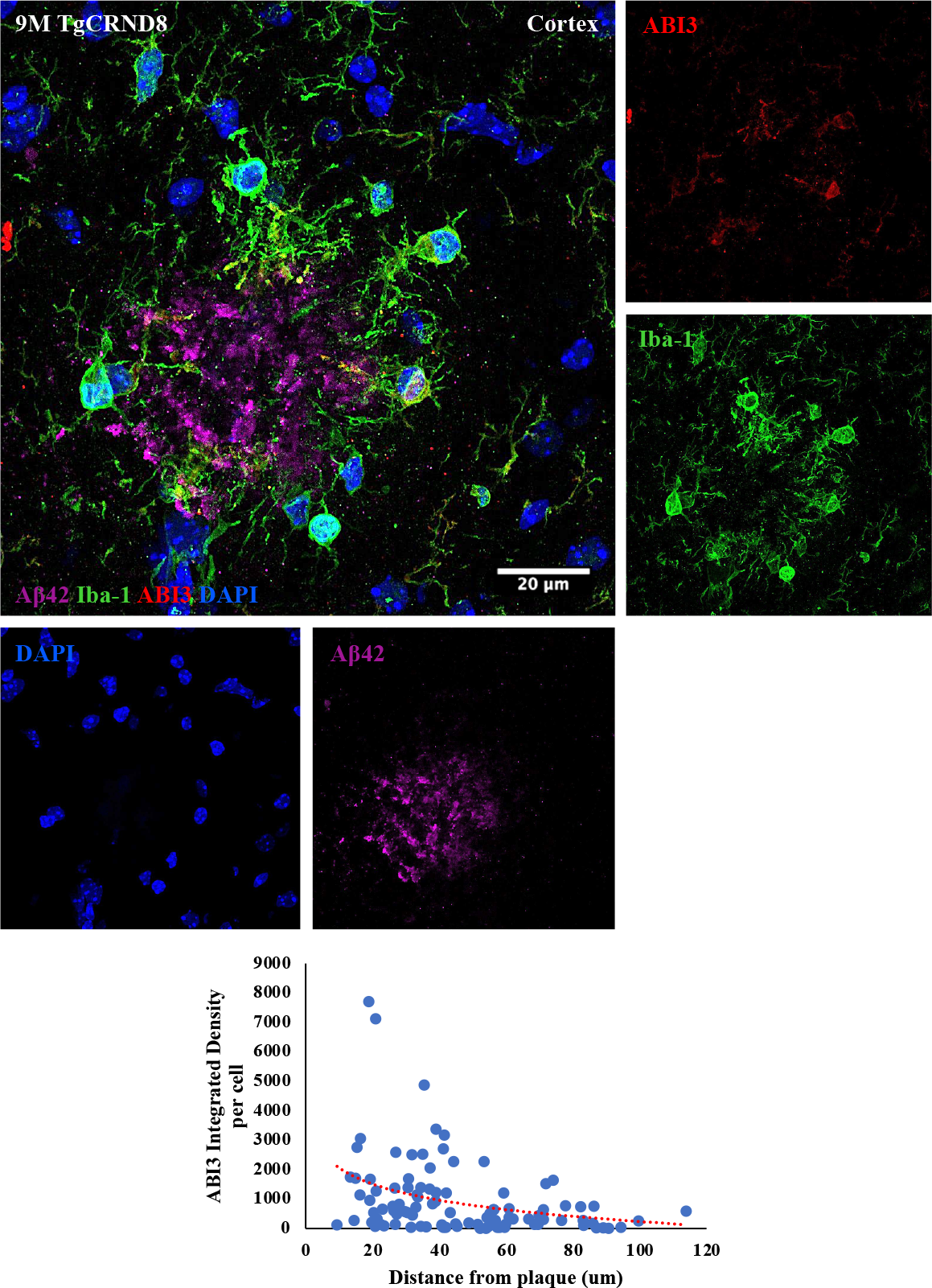
Increased ABI3 levels in TgCRND8 mice might be correlated with the microglia distance from the plaques. A representative image of Iba-1 positive microglia (green) surrounding the amyloid beta plaque (magenta) in the cortex of 9-month-old TgCRND8 mouse brain tissue. ABI3 staining (red) was observed in the microglia surrounding the plaque. The results were plotted onto a scatter graph and a moderate negative correlation was found between the cell-specific ABI3 immunoreactivity and the distance from the plaque using the Pearson correlation coefficient test (∼0.3229). N=8 TgCRND8 mice (100 measurements were taken from 16 tissue sections).

We quantified and compared several aspects of amyloid beta plaques in 9-month-old TgCRND8 ABI3-GNGT2^+/+^ (Tg WT) and Tg ABI3-GNGT2^−/−^ (Tg KO) mice. The features examined include: plaque numbers (the mean number of amyloid plaques measured in the cortex and hippocampus), plaque size (the mean Aβ42 positive pixels area (in μm2) per each plaque in the cortex and hippocampus), plaque density (%) (the percentage of mean Aβ42 positive pixels area (in μm2) divided by the whole brain area measured in each observation) and Aβ42 positive brain area (the Aβ42 positive pixels area (in μm2) measured in the cortex and hippocampus). We observed significant differences in the amyloid plaque properties in Tg +/-ABI3 WT and Tg +/-ABI3 KO mouse brains. The mean number of amyloid beta plaques measured was significantly lower in both the cortex (P value ∼ 0.0011) and the hippocampus (P value <0.0001) of Tg KO mouse brains compared to age-matched Tg WT mouse brains (**Figure 4.A**]. The amyloid plaque size, measured in μm^2^, was also significantly smaller in Tg KO mouse brains compared to age-matched Tg WT brains (P value <0.0001) (**Figure 4.B**]. Amyloid beta plaque density and the Aβ42 positive brain area were also studied in parallel as a method of providing an overview on amyloid beta burden in Tg KO and Tg WT mouse brains. Both the plaque density (%) and the Aβ42 positive brain area (in μm2) were significantly lower in Tg KO mouse brains compared to Tg WT mouse brains (P value <0.0001) [**Figure 4.C, D**].

**Figure 4:**
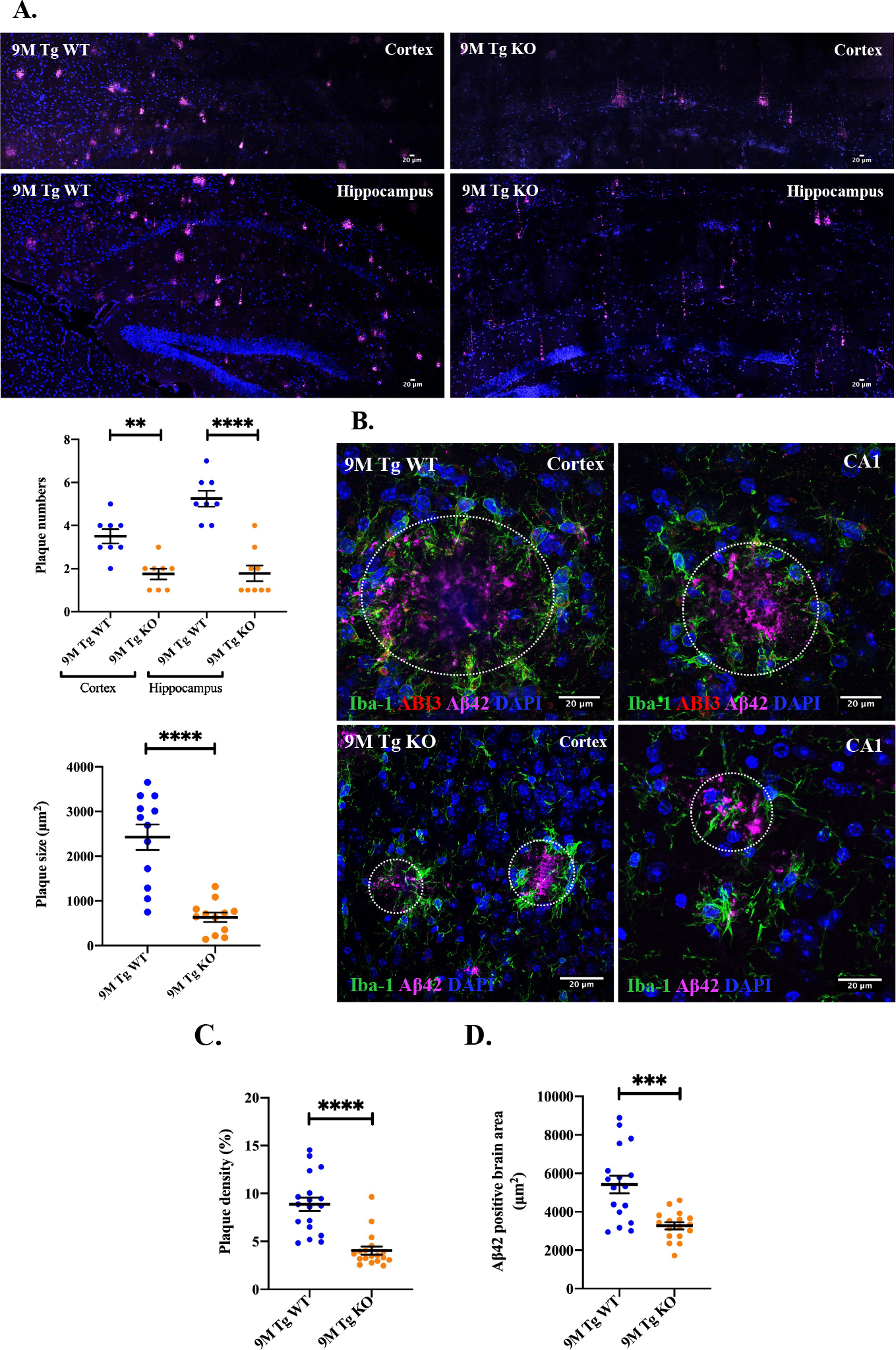
Several features of amyloid plaque pathology are attenuated in Tg KO mice. **A**. Representative images from the cortex (top) and hippocampus (CA1, bottom) of Tg WT (left) and Tg KO (right) mouse brains show reduced numbers of amyloid beta plaques (Aβ42, magenta) in Tg KO. The mean number of plaques in both the cortex (P value ∼ 0.0011) and hippocampus (P value <0.0001) of the Tg KO mouse brain was significantly lower compared to Tg WT. **B**. Representative images of amyloid plaques (magenta) and Iba-1 positive microglia (green) surrounding the plaques in the cortex (left) and hippocampus (CA1, right) of 9-month-old Tg WT (top) and Tg KO (bottom) mouse brains. The mean plaque area measured in Tg KO mouse brains was significantly lower compared to Tg WT (P value <0.0001). The plaque area is marked by the white circles. **C**. The plaque density was significantly lower in Tg KO mouse brains compared to Tg WT (P value <0.0001) and **D**. The mean Aβ42 positive brain area (measured in μm2) was significantly lower in Tg KO mouse brains compared to Tg WT. The statistical significance was obtained using Welch’s t-test. N=3 (Tg WT), n=2 (Tg KO).

We quantified and compared microglia numbers in the proximity (20μm from plaque edge) of the amyloid plaques in 9-month-old Tg WT and Tg KO mouse brains. Previous studies have reported higher microglia numbers within the radius of 20-30μm from the plaque edge and significant synapse loss within the distance of 20μm from the plaque edge^10^. Hence the 20μm radius from the plaque edge was chosen as the critical cut-off value. The mean number of microglia per plaque was significantly higher in the 20μm radius from the plaque edge in both Tg WT (P value <0.0001) and Tg KO mouse brains (P value ∼0.0001). The mean number of microglia in the proximity of plaques (20μm radius from the plaque edge) was significantly lower in Tg KO mouse brains compared to Tg WT mice (P value∼0.0021) [**Figure 5**].

**Figure 5:**
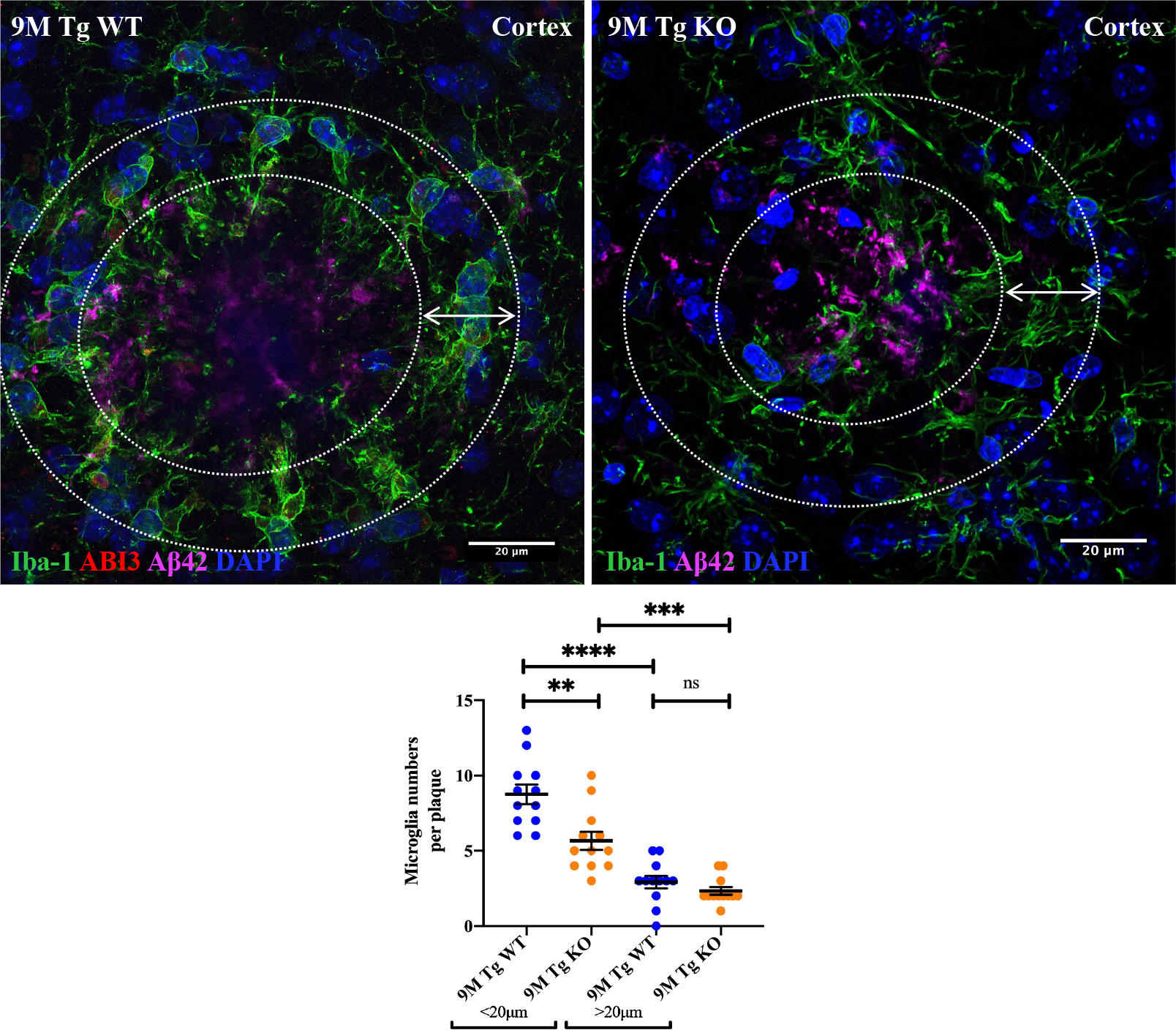
Reduced microglial clustering around the plaques in Tg KO mice. Representative images showing the Iba-1 positive microglia (green) associated with amyloid plaques (magenta) in the cortex of 9-month-old Tg WT (left) and age-matched Tg KO (right) mouse brains. The 20μm radius of the plaque edge is marked with white circles and arrows. The bottom graph shows the statistical analysis of the mean number of microglia surrounding each amyloid plaque in Tg WT and Tg KO mouse brains. Significantly higher numbers of microglia were seen in the 20μm radius of the plaque edge in both Tg WT (P value <0.0001) and Tg KO mouse brains (P value ∼0.0001), whereas a significantly lower number of microglia per each plaque was observed in the 20μm radius of the plaque edge in Tg KO mouse brains compared to the same region in the Tg WT mice (P value∼0.0021). The statistical significance was obtained using the unpaired t-test with Welch’s correction. N=3 (Tg WT), n=2 (Tg KO).

The exact mechanisms InIch, the S209F mutaIion in ABI3 increases AD risk remains unknown. ABI3 expression is highly enriched in microglia/macrophages but also observed in other cell types including neurons, thus ABI3 might have cell-specific functions and modulate AD risk through different mechanisms such as regulating tau processing as mentioned by other groups^7^. Here we characterized amyloid pathology and microglia clustering around the amyloid plaques in 9-month-old TgCRN8 ABI3-GNGT2^−/−^ mice and showed that the double deletion of ABI3-GNGT2 significantly attenuates amyloid beta pathology in these mice. Since GNGT2 transcript is also present in microglia, we cannot reject the possibility that some of the effects of ABI3-GNGT2^−/−^ could be due to loss of function of GNGT2. However, we have been unable to detect GNGT2 protein in microglia suggesting that the expression level is likely below the detection limits of assays used.

Several factors could have been attributed to the lower plaque load in TgCRND8 ABI3-GNGT2^−/−^ mice. It should first be noted that these studies were performed in 9-month-old mice and detailed time course series studies starting in younger mice (3-month-old) should be conducted in order to properly analyze the possible differences in the severity of AD pathology in TgCRN8 ABI3-GNGT2^+/+^ and ^−/−^ mice. The lower plaque numbers in TgCRND8 ABI3-GNGT2^−/−^ could have resulted from alterations in the course of disease pathogenesis and plaque formation or from improvements in the microglial ability to clear out the plaques. Given the involvement of ABI3 in actin cytoskeleton reorganization and cell movement in microglia, the latter possibility seems more plausible. A similar explanation could also be given to the average plaque size in TgCRND8 ABI3-GNGT2^−/−^ mice. In order to study this further, a multiphoton microscopy-based in vivo imaging of tagged microglia in APP mouse brains could be performed in a time-course fashion as suggested by other groups^11^.

We also recognize that opposing observations have been reported by some groups using 5xFAD ABI3-GNGT2^−/−^ mice bred on an SJL genetic background, where the AD amyloid-beta pathology is significantly increased^12^. It’s important to note that genetic background effects such as possible mutations in the genes that encode proteins upstream of ABI3 could impact ABI3 signaling (or lack of) in microglia and lead to different observations.

We also observed a significant reduction of microglia numbers around the amyloid beta plaques in TgCRN8 ABI3-GNGT2^−/−^, especially within the 20 μm radius. This could be due to different reasons. As mentioned before, either fewer plaques were formed in the first place due to loss of ABI3, which led to minor microglial activation and accumulation of microglia around the plaques; or a higher number of microglia had migrated to the sites of amyloid plaques and were better able to clear out the plaques, which by the time of the analysis a fewer number of microglia seemed to be associated with the plaques. If we associate microglial accumulation around amyloid plaques as one of the hallmarks of “gliosis” in AD, fewer plaque-associated microglia in TgCRND8 ABI3-GNGT2^−/−^ mice could suggest a possible involvement of ABI3 in microglial response in AD. However, more extensive research should be performed by analyzing different aspects of microglial response in AD, at different time points. These observations would then provide a better understanding of ABI3 involvement in microglial functions in AD and the risk effects of S209F mutation.

## Methods

### Antibodies

The following antibodies were used: GAPDH (# 2118, Cell Signaling), β-Amyloid 1-16 (# 803014, BioLegend), ABI3 (# ab81152, abcam), Iba1 (# NB100-1028, Novus Biologicals). Donkey anti-Goat IgG (H+L) Cross-Adsorbed Secondary Antibody, Alexa Fluor™ 488 (# A-11055, ThermoFisher), Donkey anti-Mouse IgG (H+L) Highly Cross-Adsorbed Secondary Antibody, Alexa Fluor™ 647 (# A-31571, ThermoFisher), Donkey anti-Rabbit IgG (H+L) Highly Cross-Adsorbed Secondary Antibody, Alexa Fluor™ 594 (# A-21207, ThermoFisher).

### Animals

All animal studies were approved by the Animal Care Committees of the University of Toronto and the University Health Network and were performed in accordance with the guidelines of the Canadian Council on Animal Care. Abi3tm1.1(KOMP)Vlcg (Abi3-deficient) mice were purchased from the Jackson Lab. This strain was created using a targeted deletion strategy with a reporter-tagged cassette, later removed by cre. It has a 9515bp deletion eliminating exons 1-8 are maintained on a C57/Bl6 background. Abi3-deficient animals are maintained as KO X KO animals on a C57/Bl6 background. For neuropathological studies, Abi3-deficient animals were cross-bred with the TgCRND8 APP transgenic mouse (C57:C3H background). A two-stage breeding strategy was used to create animals that were hemizygous for the human APP transgene and homozygous for the Abi3 gene ablation: APP^+/-^ ABI3^−/−^. The presence of the APP (TgCRND8 strain; PMID: 11279122) was confirmed by genomic DNA dot blot.

### Brain extraction and preparation

Brain tissue was collected from animals at 9 months of age. Briefly, mice were terminally anesthetized with sodium pentobarbital (200mg/kg), on reaching the anesthetic plane they were transcardially perfused with 30ml of PBS, brains were then removed and fixed in 4% phosphate-buffered paraformaldehyde for 24h then transferred to 30% sucrose for cryoprotection. The brains were embedded in OCT compound then snap-frozen on dry ice for in preparation for cryosectioning. X micron thick sections were placed on Y slides.

### Histopathological studies

Frozen brain sections were briefly dehydrated using methanol and antigen retrieval was performed with 1X Antigen Retrieval Buffer (100X Citrate Buffer) (ab93678). Slides were blocked in 10% Donkey Serum in 1X TRIS_buffered saline containing 0.3% Triton X-100 (TBS-T)for 1h at room temperature followed by primary antibody incubation at 4 °C overnight. Appropriate secondary antibodies were used for blocking at room temperature for 1h. Slides were mounted in ProLong™ Diamond Antifade Mountant with DAPI (# P36962, ThermoFisher).

### Image acquisition, analysis and statistics

Zeiss LSM880 confocal microscope was used for image acquisition. For each brain tissue, two sections were imaged and quantified. Quantifications were performed using ImageJ (Fiji) software (Version 1.0) and all statistical analyses were performed using Prism (Version 8.0 GraphPad Software Inc.). Multiple comparisons used Kruskal-Wallis analysis of variance methods (ANOVA), and ANOVA with Tukey multiple comparisons post-test. The specific statistical tests used in each experiment are described in the associated figure legend. Data are expressed as means ± SEM and results were considered significant if p<0.05.

## Acknowledgments

This work was supported by grants from Canadian Institutes of Health Research (406915 Foundation Grant and Canadian Consortium on Neurodegeneration in Aging Grant), US Alzheimer Society Zenith Grant ZEN-18-529769, Alzheimer Society of Ontario Chair in Alzheimer’s Disease Research and National Institute of Aging (U01AG072572; R01AG070864).

## Author contributions

PHStGH designed the study. DG performed the experiments, analyzed the data, and wrote the manuscript. JG provided the mice and processed the tissue sections.

